# Aspirin reprogrammes colorectal cancer cell metabolism and sensitises to glutaminase inhibition

**DOI:** 10.1101/2022.08.24.505115

**Authors:** Amy K. Holt, Arafath K. Najumudeen, Ashley J. Hoskin, Danny N. Legge, Eleanor M.H. Mortensson, Dustin J. Flanagan, Nicholas Jones, Madhu Kollareddy, Tracey J. Collard, Penny Timms, Owen J. Sansom, Ann C. Williams, Emma E. Vincent

## Abstract

To support proliferation and survival within a challenging microenvironment, cancer cells must reprogramme their metabolism. As such, targeting cancer cell metabolism is a promising therapeutic avenue. However, identifying tractable nodes of metabolic vulnerability in cancer cells is challenging due to their metabolic plasticity. Identification of effective treatment combinations to counter this is an active area of research. Aspirin has a well-established role in cancer prevention, particularly in colorectal cancer (CRC), although the mechanisms are not fully understood. Here, we comprehensively characterise the metabolic impact of long-term aspirin exposure (2-4mM for 52 weeks) on CRC cells. We show that aspirin regulates several enzymes and transporters of central carbon metabolism and results in a reduction in glutaminolysis and a concomitant increase in glucose metabolism, demonstrating reprogramming of nutrient utilisation. We show that aspirin causes likely compensatory changes that renders the cells sensitive to the glutaminase 1 (GLS1) inhibitor - CB-839. Of note given the clinical interest, treatment with CB-839 alone had little effect on CRC cell growth or survival. However, in combination with aspirin, CB-839 inhibited CRC cell proliferation and induced apoptosis *in vitro*, and importantly, reduced crypt proliferation in *Apc^fl/fl^* mice *in vivo*. Together, these results show that aspirin leads to significant metabolic reprogramming in colorectal cancer cells and raises the possibility that aspirin could significantly increase the efficacy of metabolic cancer therapies in CRC.

## Introduction

Metabolic reprogramming is a defining feature of cancer cells (1) and is essential to meet both the energetic and biosynthetic requirements of chronic proliferation. Although the specific nature of cancer cell metabolism depends on several factors including tissue of origin and mutational status, often common features include increased aerobic glycolysis (known as the Warburg effect) and glutamine utilisation (2, 3).

Colorectal cancer (CRC) is the second most common cause of cancer related death in the UK, and incidence is increasing, particularly in younger patient populations (4, 5), highlighting the need for improved therapies. Metabolic reprogramming supports CRC initiation and progression and it is well established that it is a driver rather than a passive outcome of tumourigenesis. Indeed, many key oncogenic pathways in CRC have been shown to directly control metabolism, including Wnt, PI3K and p53 signalling (6, 7). As such, cancer metabolism is an attractive target for novel therapies. However, many challenges remain in developing metabolic anti-cancer therapies, such as the large overlap between the metabolic programme favoured by cancer cells and normal proliferating cells, resulting in a small therapeutic window and the increased likelihood of toxicity (8). There is also the challenge of overcoming metabolic plasticity; cancer cells are well suited to adapting their metabolism to meet environmental constraints, such as hypoxia and hypoglycaemia, and the contrasting conditions of the bloodstream and metastatic sites (9, 10). As a result, when targeted with singular metabolic interventions, cancer cells often rewire their metabolism to enable continued proliferation.

Aspirin is a widely used non-steroidal anti-inflammatory drug (NSAID) prescribed for the prevention of cardiovascular events in high-risk patients. A growing body of epidemiological evidence suggests that aspirin reduces cancer incidence, in particular CRC, as well as potentially slowing disease progression (11, 12). The US Preventative Services Task Force recommend daily aspirin for CRC prevention in 50–69 year-olds with increased risk of cardiovascular disease and no increased risk of bleeding (13). Furthermore, the National Institute for Health and Care Excellence (NICE) guidelines now recommend daily aspirin for the prevention of CRC in patients with Lynch syndrome (14).

Aspirin is a pleiotropic drug; its actions at the cellular level are not fully understood, particularly with regards to its role in cancer prevention. Increased knowledge of aspirin’s cellular mechanisms could enhance its efficacy, including identification of optimal timing and dose, and those individuals most likely to benefit from taking regular aspirin.

Epidemiological data suggest that the effect of aspirin on CRC incidence and progression is affected by the length of time for which aspirin is taken (11, 15). While the effects of aspirin have been extensively studied *in vitro* and *in vivo*, long-term exposure has not been modelled before in cell lines. Therefore, in this study, we investigated the impact of long-term (52 week) aspirin exposure on CRC cells, with the aim of identifying novel mechanisms of action. Detailed proteomic and metabolomic analysis revealed altered metabolism and nutrient utilisation with aspirin exposure.

Several key enzymes involved in central carbon metabolism were identified as being regulated by aspirin, including pyruvate carboxylase (PC), pyruvate dehydrogenase kinase 1 (PDK1) and glutaminase 1 (GLS1). Although aspirin alone did not impact the ability of the cells to produce ATP, it does inhibit net glutaminolysis, despite inducing a (likely compensatory) increase in GLS1 expression. Importantly, although the GLS1 inhibitor CB-839 alone had little effect on CRC cell survival, aspirin renders colorectal cells sensitive to the drug both *in vitro* and *in vivo*. In addition, reduced glutaminolysis upon aspirin exposure leads to a concomitant and likely compensatory increase in glucose utilisation in the tricarboxylic acid (TCA) cycle, leaving cells sensitive to the mitochondrial pyruvate carrier 1 (MPC1) inhibitor, UK-5099. In summary, we demonstrate that aspirin causes metabolic rewiring in colorectal cancer cells providing therapeutic opportunities to sensitise colorectal cancer to existing metabolic cancer therapies currently under clinical investigation.

## Results

### Long-term aspirin exposure regulates expression of metabolic pathway genes in CRC cells

To explore the consequences of long-term aspirin exposure on SW620 colorectal cancer cells, we performed proteomic analysis to compare protein expression in cells treated for 52 weeks in continuous culture with either 2mM or 4mM aspirin to untreated controls (experimental design shown in Figure 1a). 265 proteins were significantly differentially regulated in cells treated with both 2mM and 4mM aspirin compared to untreated controls (p<0.05, fold change>1.4 or <0.71) (16) (Figure 1b). Analysis of these proteins using Webgestalt highlighted “metabolic process” as having the highest number of genes in the gene ontology (GO) biological processes (Figure 1c). Overrepresentation analysis using the KEGG pathway database highlighted a high enrichment ratio in “metabolic pathways” and “central carbon metabolism in cancer”, as well as some specific metabolic pathways including “pyruvate metabolism”, “cholesterol metabolism” and “fatty acid biosynthesis” (Figure 1d). These results suggest that long-term aspirin exposure might rewire cellular metabolic pathways in CRC cells.

**Figure 1.**
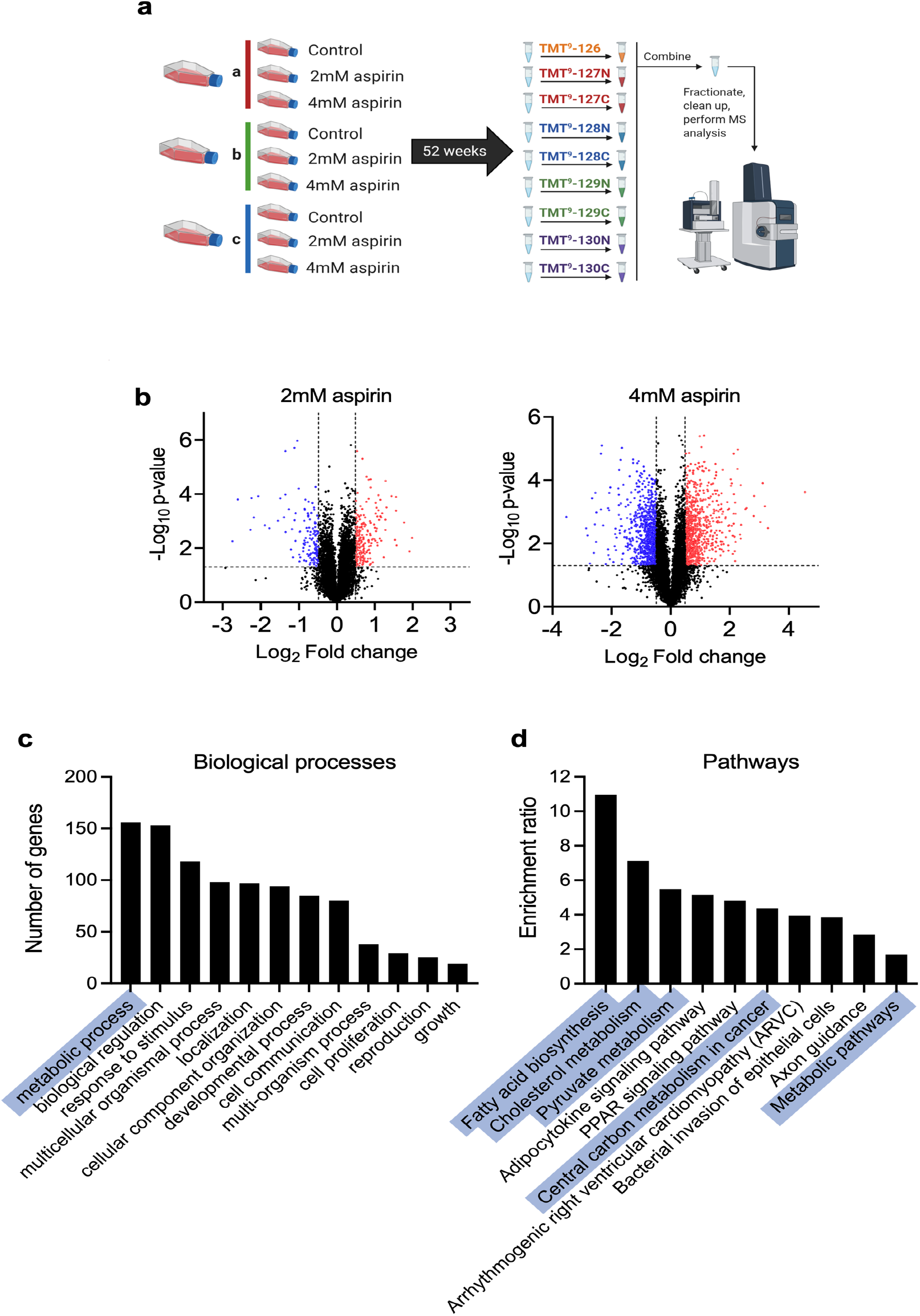
Long-term aspirin treatment regulates cellular metabolism in CRC cells. **a)** Experimental design for proteomic analysis of long-term aspirin treated SW620 cells (each flask represents one independent experiment) (created with BioRender.com). **b)** Volcano plots showing protein expression changes with long-term 2mM or 4mM aspirin treatment compared to control. Each point represents one protein. Thresholds for proteins of interest were p<0.05, fold change>1.4 or <0.71), in both 2mM and 4mM conditions (downregulated proteins in in blue and upregulated in red). **c)** Number of genes in GO biological processes categories, of all proteins of interest in both 2mM and 4mM aspirin. **d)** Overrepresentation analysis of proteins of interest in both 2mM and 4mM aspirin, using KEGG pathways database.

### Aspirin exposure reprogrammes nutrient utilisation in CRC cells

To determine whether aspirin exposure leads to a functional change in energy metabolism as predicted by the proteomic analysis, we next investigated the effect of long-term aspirin exposure on ATP production from glycolysis and oxidative phosphorylation (oxphos) using a Seahorse Extracellular Flux Analyzer. Surprisingly, no changes were observed in either basal or maximal oxygen consumption rate (OCR) or extracellular acidification rate (ECAR), proxy measures of oxidative and glycolytic activity respectively, with long-term aspirin treatment (Figure 2a). This suggests that aspirin exposure has no net impact on ATP production in CRC cells and is therefore unlikely to explain the known effect of aspirin on cellular proliferation (17).

**Figure 2.**
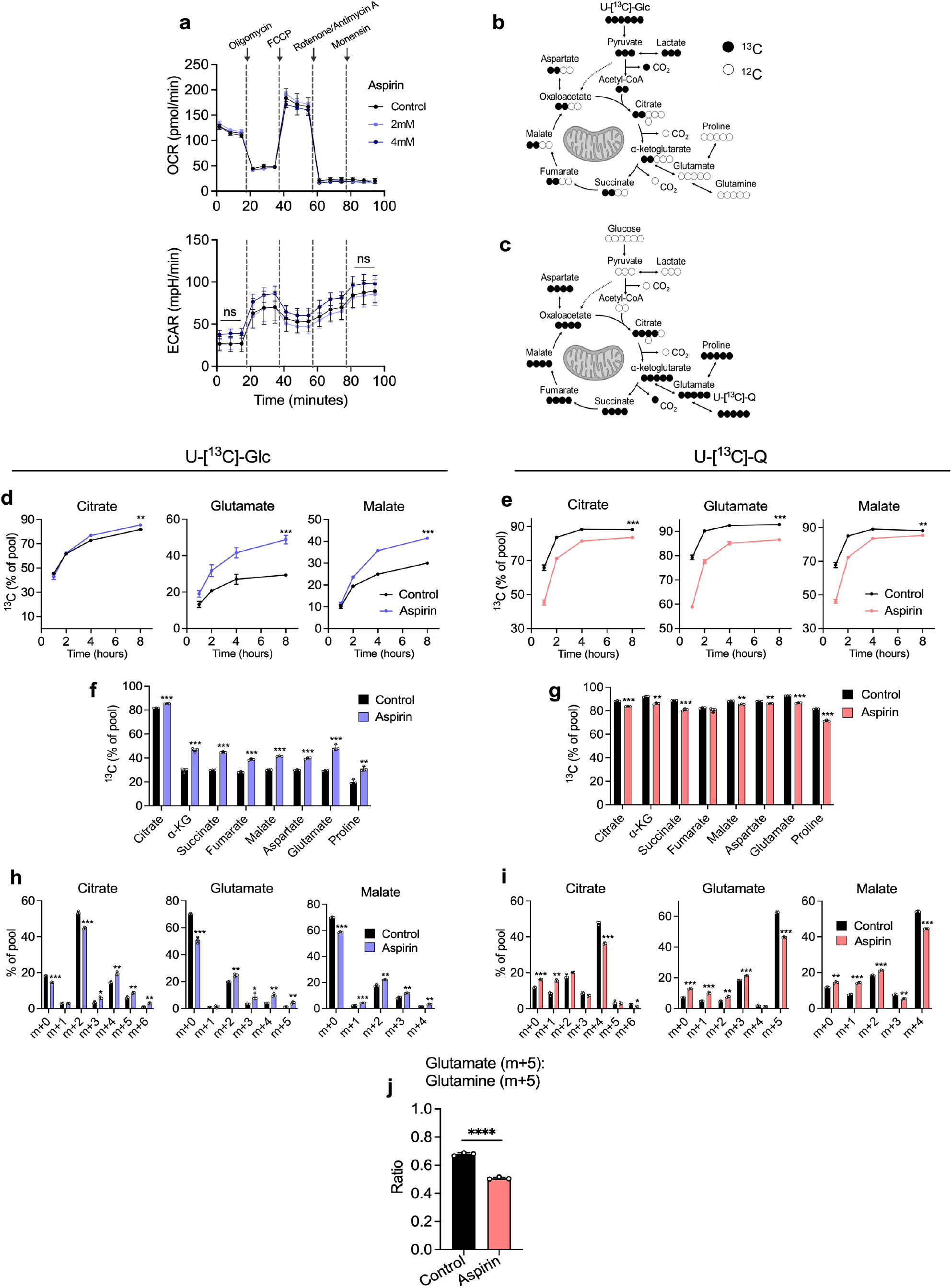
Long-term aspirin treatment reprogrammes nutrient utilisation in CRC cells. **a)** Extracellular flux analysis of long-term (52 week) 2mM and 4mM aspirin treated SW620 cells compared to controls. Error bars represent SEM (n=3 independent experiments). ns=not significant (p>0.05 following t tests at indicated time points). **b-c)** Schematics of U-[^13^C]-Glc (b) and U-[^13^C]-Q (c) incorporation into TCA cycle metabolites and amino acids. Created with BioRender.com. **d-j)** SIL data for long-term (52 week) 4mM aspirin treated SW620 cells compared to control cells. Error bars represent SD (n=3 technical replicates). Asterisks refer to p-values obtained from t tests at the 8 hour time point (*=p<0.05, **=p<0.01, ***=p<0.001, ****=p<0.0001). **d-e)** Proportion of ^13^C labelling in citrate, glutamate and malate from U-[^13^C]-Glc and U-[^13^C]-Q over time. **f-g)** Proportion of ^13^C labelling in metabolite pools at 8 hours from U-[^13^C]-Glc (f) and U-[^13^C]-Q (g). Asterisks indicate the adjusted p-value obtained using multiple t-tests. **h-i)**Mass Isotopomer Distribution (MID) analysis at 8 hours for U-[^13^C]-Glc (h) and U-[^13^C]-Q (i) labelling in citrate, glutamate and malate. Asterisks indicate the adjusted p-value obtained using multiple t-tests. **j)**Ratio of m+5 glutamateo m+5 glutamine in long-term 4mM aspirin treated cells in comparison to control at 8 hours from U-[^13^C]-Q.

We next conducted stable isotope labelling (SIL) experiments to investigate whether aspirin altered the metabolic fate of glucose and glutamine, the most important carbon sources for proliferating cancer cells in culture. Long-term (52 week) aspirin treated SW620 cells were incubated with either uniformly labelled ^13^C-glucose (U-[^13^C]-Glc) or glutamine (U-[^13^C]-Q) for up to 8 hours in order to capture isotopic steady state (18). Conventional metabolism of U-[^13^C]-Glc and U-[^13^C]-Q in tumour cells is illustrated in Figure 2b-c. The majority of metabolites reach isotopic steady state within 8 hours of incubation (shown for citrate, glutamate and malate), with the steady state proportion of labelled metabolite (indicating labelled nutrient contribution) being altered by aspirin exposure (Figure 2d-e). Incorporation of U-[^13^C]-Glc and U-[^13^C]-Q across TCA cycle metabolites at 8 hours was increased and decreased respectively upon aspirin exposure (Figure 2f-g). Similar results were observed in LS174T and HCA7 cells exposed to long-term aspirin (Supplementary Figure 1).

Mass isotopomer distribution (MID) analysis of citrate, glutamate and malate shows a decrease in the unlabelled metabolites (m+0) and an increase in the proportion of the glucose labelled mass isotopomers with aspirin exposure (Figure 2h). By contrast, an increase in unlabelled metabolites (m+0) and a decrease in the proportion of the glutamine labelled isotopomers was observed (Figure 2i). These data suggest that glutaminolysis is inhibited by aspirin exposure, confirmed by analysis of the glutamate:glutamine m+5 ratio (Figure 2j).

These data demonstrate that despite there being no overall impact of long-term aspirin exposure on ATP production, it does cause metabolic reprogramming in three different CRC cell lines, reducing glutaminolysis and increasing glucose utilisation. Glucose and glutamine cooperate in fuelling the TCA cycle; decrease in entry of one nutrient can lead to a compensatory increase in the other (19, 20). This suggests that the increased glucose utilisation may be a compensation mechanism for the reduction in glutaminolysis in the presence of aspirin in order to maintain carbon entry into the TCA cycle. As there was no overall effect on oxphos, this suggests the cells can maintain TCA cycle function in the presence of aspirin by increasing glucose utilisation. Taken together, these data highlight the metabolic plasticity of the cells, allowing them to minimise the impact on ATP production in the presence of aspirin.

### Aspirin regulates levels of proteins involved in central carbon metabolism

Having determined that aspirin impacts cellular metabolism in CRC cells, we sought to identify changes in protein expression that are consistent with the metabolic rewiring we observe. For this we performed further analysis on the proteomic data in Figure 1 and found that Ingenuity Pathway Analysis (IPA) predicted inhibition of Activating Transcription Factor 4 (ATF4, an important regulator of cellular metabolism) signalling with aspirin, (p= 0.0116, z-score=−2.894). Regulation of all ATF4 target genes captured by IPA are shown in Figure 3a.

**Figure 3.**
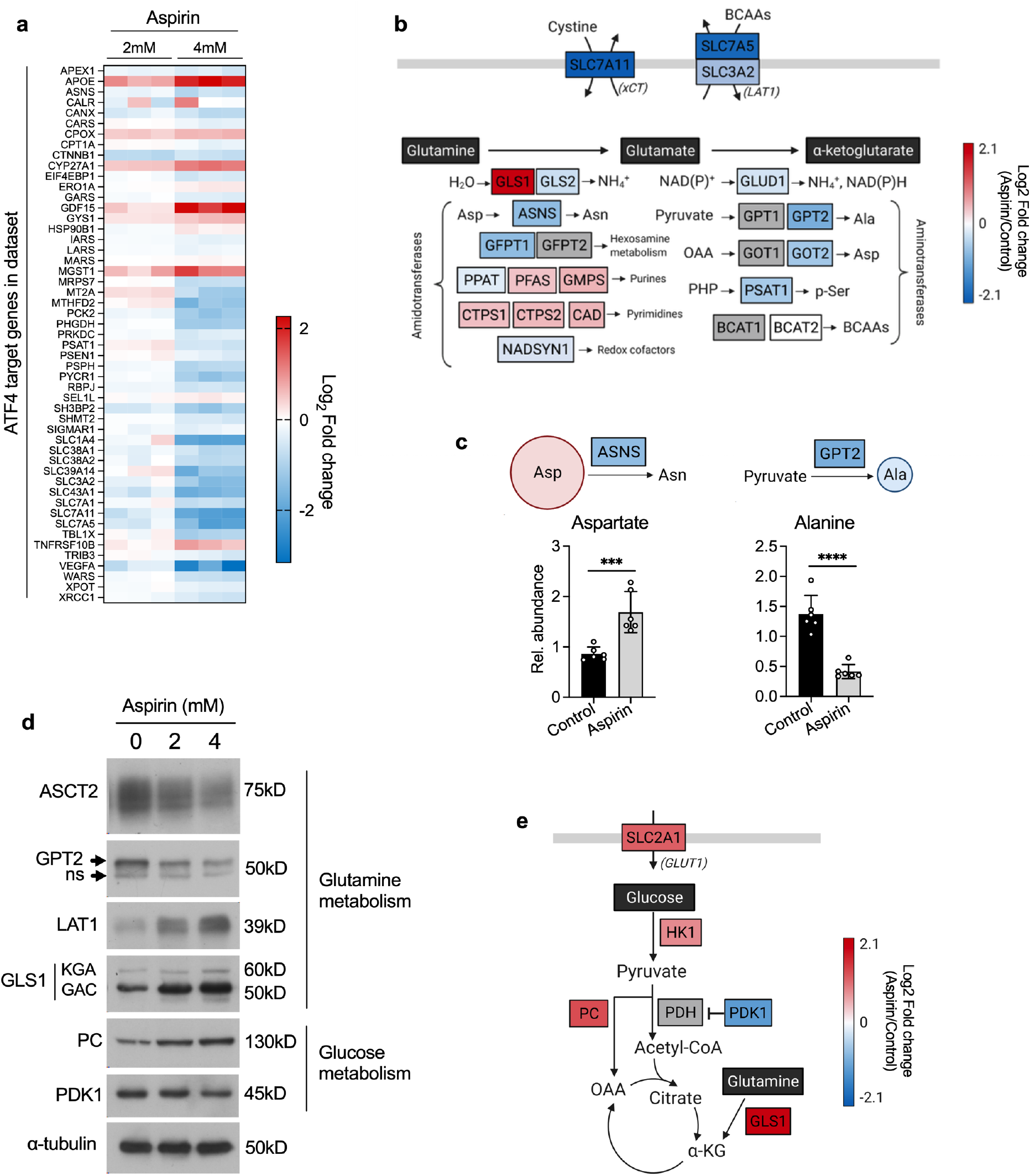
Aspirin regulates proteins involved in central carbon metabolism. **a)** ATF4 target genes highlighted in proteomic data by IPA analysis. Fold change of proteins in both long-term (52 week) 2mM and 4mM aspirin conditions, relative to control (n=3 technical replicates). IPA analysis shows a predicted overall inhibition of ATF4 signalling with long-term 4mM aspirin (p= 0.0116, z-score=−2.894). **b)** Overview of key enzymes involved in glutaminolysis and their average fold changes with long-term 4mM aspirin treatment in the proteomic data. BCAAs=branched chain amino acids, OAA=oxaloacetate, PHP=phosphohydroxypyruvate. Created with BioRender.com. **c)** Metabolite abundance relative to cell number of alanine and aspartate in long-term 4mM aspirin treated cells compared to control. Error bars represent SD (n=6 technical replicates). Asterisks refer to p-values obtained using t tests ***=p<0.001, ****=p<0.0001). Created with BioRender.com. **d)** Immunoblotting for a selection of metabolic enzymes highlighted in proteomic data, with long-term aspirin treatment. Representative of at least 3 independent experiments. α-tubulin is used as a loading control. **e)**Average fold changes in the proteomic data with long-term 4mM aspirin compared to control, including central carbon metabolism genes that showed significant regulation (p<0.05, fold change>1.4) in both long-term 2mM and 4mM aspirin. Created with BioRender.com.

ATF4 is involved in the cellular response to amino acid deprivation and has also been shown to regulate glutamine metabolism (21, 22). Figure 3b shows an overview of proteins involved in glutamine metabolism and transport, several of which are ATF4 target genes. For example, expression of glutamic-pyruvic transaminase 2 (GPT2), which catalyses the reversible transamination between alanine and a-ketoglutarate (aKG) to generate pyruvate and glutamate was downregulated upon aspirin treatment (4mM aspirin vs control; log_2_ fold change = −0.95, p-value = 0.03) and has been previously shown to be regulated by ATF4 (22). There was also downregulation of ATF4 targets; asparagine synthetase (ASNS; log_2_ fold change= −0.75, p-value= 0.004) and phosphoserine aminotransferase 1 (PSAT1; log_2_ fold change= −0.58, p-value = 0.0008), as well as glutamate dehydrogenase 1 (GLUD1; log_2_ fold change= −0.27, p-value= 0.013) and the aminotransferase glutamic-oxaloacetic transaminase 2 (GOT2; log_2_ fold change= −0.62, p-value= 0.004). Furthermore, two amino acid transporters that impact glutamine metabolism; cystine/glutamate antiporter (xCT, encoded by the *SLC7A11* gene; log_2_ fold change= −2.09, p-value= 0.0009) and the large neutral amino acid transporter LAT1 (encoded by the *SLC7A5* gene; log_2_ fold change= −1.89, p-value= 0.012), also highlighted by the IPA in Figure 3a, were both downregulated with aspirin. These data are consistent with the reduced levels of glutaminolysis we observed in Figure 2. Also consistent is an increase in intracellular abundance of aspartate and decrease in intracellular abundance of alanine upon long-term aspirin exposure, potentially illustrating the functional consequence of reduced expression of ASNS and GPT2 respectively (Figure 3c).

Consistent with the proteomic data, mRNA levels of *GPT2, SLC7A11* and *SLC7A5*, were found to be downregulated upon long-term aspirin exposure, suggesting strong transcriptional regulation (Supplementary Figure 2a). Neither protein nor mRNA levels of ATF4 itself were significantly regulated with long-term aspirin exposure (Supplementary Figure 2b), consistent with post-transcriptional regulation of ATF4 (23). Validation of proteomic data was performed for GPT2 and LAT1 by immunoblotting (Figure 3d and Supplementary Figure 2c). The key glutamine transporter ASCT2 (alanine-serine-cysteine transporter 2) was also investigated by immunoblotting and was downregulated with long-term 4mM aspirin (Figure 3d), although this was not statistically significantly (Supplementary Figure 2c).

Further analysis of the proteomic data also revealed expression changes consistent with the concomitant increase in glucose metabolism we observed in Figure 2f. Two proteins involved in regulating entry of pyruvate into the TCA cycle showed significant regulation – PC was upregulated (log_2_ fold change= +1.28, p-value= 0.00037) and PDK1 was downregulated (log_2_ fold change= −1.19, p-value= 0.002) (illustrated in Figure 3e). These results were validated by immunoblotting (Figure 3d and Supplementary Figure 2d. This is consistent with the increased glucose carbon entry into the TCA cycle that we previously observed (Figure 2f and h). In addition, glycolysis enzyme hexokinase 1 (HK1; log_2_ fold change=+0.79, p-value= 0.012) and glucose transporter 1 (GLUT1, encoded by the *SLC2A1* gene; log_2_ fold change=+1.13, p-value= 0.00026) were both upregulated in the proteomic data, also consistent with increased glucose utilisation upon aspirin exposure (Figure 3e).

Unexpectedly, GLS1, which catalyses the first step of glutaminolysis (Figure 3b), showed significant upregulation (Figure 3d-e and Supplementary Figure 2d). This is inconsistent with the reduced levels of glutaminolysis we observed in Figure 2. Both known splice variants of GLS1 (GLS1^KGA^ and GLS1^GAC^) were identified (Figure 3d), with GLS1^GAC^ being the dominantly expressed isoform in our cells. qPCR analysis of aspirin treated SW620 cells did not show any significant transcriptional regulation of *GLS1, PC* or *PDK1* (Supplementary Figure 2e), suggesting post-transcriptional regulation of these proteins.

To demonstrate the changes in other CRC cell lines, expression of GLS1, PC and PDK1 was also investigated by immunoblotting in LS174T and HCA7 cells after long-term aspirin exposure (Supplementary Figure 3a), showing upregulation of GLS1^GAC^ in LS174T, though this is not statistically significant, and significant downregulation of PC and PDK1 in HCA7 cells.

Expression changes were also investigated by immunoblotting following short-term aspirin treatment (72 hours) (Supplementary Figures 3a-b). This showed significant upregulation of GLS1^GAC^ in SW620 and LS174T cells, as well as upregulation of PC in SW620. In addition, downregulation of mRNA expression of the ATF4 targets *GPT2* and *SLC7A5* was also observed in SW620 cells (Supplementary Figure 3c). Interestingly, these findings suggest short-term treatment is sufficient for the regulatory effect of aspirin on these metabolic enzymes. However, it should be noted that long-term aspirin exposure has a stronger effect.

### Metabolic reprogramming in response to aspirin exposes metabolic vulnerabilities in CRC cells

While the metabolic impact of aspirin may be insufficient to explain the known detrimental effect on cellular proliferation (17), it could render cells more susceptible to further metabolic perturbation. Despite an overall reduction in glutaminolysis (Figure 2g), aspirin causes a strong upregulation in GLS1 levels (Figure 3d), which may be a compensatory mechanism to maximise utilisation of glutamine when levels of other glutaminolysis enzymes are reduced (such as GPT2). Increased expression of GPT2 has been previously shown to compensate for inhibition of GLS1, suggesting that the reverse relationship may also occur (24). We therefore hypothesised that aspirin treated cells may be more sensitive to further blockade of glutaminolysis by targeting GLS1. To investigate this, long-term aspirin exposed cells were incubated with increasing concentrations of CB-839 (a selective GLS1 inhibitor currently in clinical trials (25), also known as Telaglenastat, illustrated in Figure 4a). SW620 cells showed no sensitivity to CB-839 alone (up to 10μM), consistent with previous findings (26), however, cells exposed to long-term aspirin showed significantly increased sensitivity in a dose-dependent manner (Figure 4b). Similar results were obtained in long-term aspirin treated HCA7 cells and to a lesser extent LS174T. Although both of these cell lines showed minimal sensitivity to CB-839 without aspirin, aspirin significantly increased their response to the drug (Supplementary Figure 4a-b). This effect was also investigated with short-term aspirin treatment in SW620 cells (Supplementary Figure 4c). This also showed sensitisation, but to a lesser extent than in long-term aspirin treated cells.

**Figure 4.**
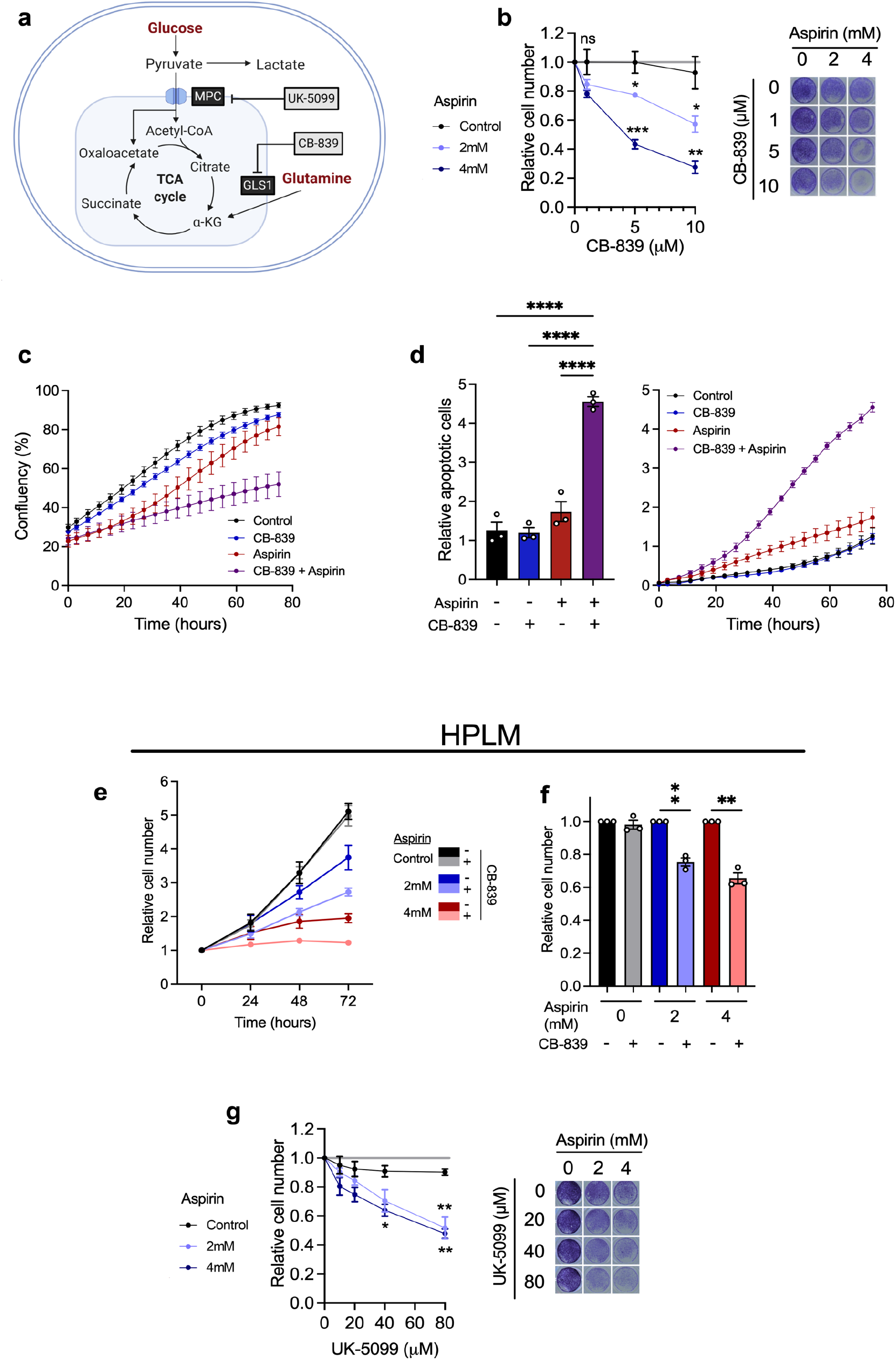
Aspirin treatment sensitises CRC cells to metabolic inhibitors. **a)** Schematic showing mechanism of action of metabolic inhibitors; CB-839 and UK-5099. Created with BioRender.com. **b)** Cell proliferation assay of long-term (52 week) aspirin treated SW620 cells with increasing concentration of CB-839. Graph shows relative cell number in each aspirin condition, measured by crystal violet staining at 72 hours compared to vehicle control. Error bars show SEM (n=3 independent experiments). Asterisks refer to p-values obtained using one-way ANOVAs with Dunnett’s multiple comparisons tests at each CB-839 concentration (*=p<0.05, **=p<0.01, ***=p<0.001). Images show representative wells in each condition at 72 hours. **c-d)** Confluency and relative apoptotic cells in long-term 4mM aspirin treated SW620 cells in combination with 5μM CB-839, in comparison to controls and to each drug alone. Error bars show SEM (n=3 distinct passages of cells analysed on the same experimental plate). Relative apoptotic cells were measured by green fluorescent nuclei (indicating cells with activated caspase-3/7) relative to cell confluency. Line graphs show values over time and bar graph shows the values at the experiment end point (75 hours after treatment). Error bars show SEM (n=3 distinct passages of cells analysed on the same experimental plate). Asterisks refer to p-values obtained using a one-way ANOVA with Tukey’s multiple comparisons test (****=p<0.0001). **e-f)** Proliferation assay of SW620 cells treated with aspirin and/or CB-839 compared to vehicle control for 72 hours, performed in Human Plasma-Like Medium (HPLM). **e)** Cell number over time relative to the 0 hour time point. **f)** Relative cell number at 72 hours, 5μM CB-839 treatment condition is shown relative to vehicle control in the same aspirin treatment condition. Error bars show SEM (n=3 independent experiments). Asterisks refer to p-values obtained using one sample t-tests, comparing to a hypothetical mean of 1 (**=p<0.01). **g)** Cell proliferation assay of long-term aspirin treated SW620 cells with increasing concentration of UK-5099. Graph shows relative cell number in each aspirin condition measured by crystal violet staining at 72 hours. Error bars show SEM (n=3 independent experiments). Asterisks refer to p-values obtained using one-way ANOVAs with Dunnett’s multiple comparisons tests at each UK-5099 concentration (*=p<0.05, **=p<0.01, ***=p<0.001). Images show representative wells in each condition at 72 hours.

To further investigate the effect of CB-839 on long-term aspirin exposed cells, proliferation assays were performed alongside detection of caspase-3/7 activation to quantify levels of apoptosis using an Incucyte® Live-Cell Analysis System (Figure 4c-d). These results confirm the inhibitory effect on proliferation of combined CB-839 and long-term aspirin, as shown by a decrease in confluency (Figure 4c). These results also show significant induction of apoptosis with the combination of CB-839 and long-term aspirin treatment compared to vehicle control and to either drug alone (Figure 4d). A positive control for this assay was performed using ABT-737 to induce apoptosis (Supplementary Figure 4d-e).

Proliferation experiments were also performed using human plasma-like medium (HPLM), developed by Cantor et al. to be representative of the metabolite composition of human plasma (27). Similar results were obtained in HPLM to those performed in DMEM (Figure 4e-f). Both 2mM and 4mM aspirin inhibited proliferation compared to controls. Addition of CB-839 in the absence of aspirin had no effect on proliferation, whereas it significantly reduced cell number with both 2mM and 4mM aspirin. This demonstrates that the effect of aspirin on sensitising cells to CB-839 is present in physiologically relevant metabolic conditions.

Upon long-term aspirin exposure, we have shown that glutaminolysis is reduced (Figure 2j), leading to a potentially compensatory increase in (Figure 3b and d), and dependence on, GLS1 (Figure 4b-f). We hypothesised that the increase in glucose utilisation we observed in Figure 2f is another compensatory response to impaired glutaminolysis in order to maintain TCA cycle activity. We reasoned this could leave cells vulnerable to inhibition of glucose utilisation and specifically to pyruvate import into the mitochondria. We investigated this by treating cells exposed to long-term aspirin with increasing concentrations of an inhibitor of the mitochondrial pyruvate carrier 1 (MPC1), UK-5099. UK-5099 inhibits entry of glucose-derived pyruvate into the TCA cycle (illustrated in Figure 4a). SW620 cells showed little or no sensitivity to UK-5099 alone, but sensitivity was significantly increased in cells exposed to both 2mM and 4mM aspirin (Figure 4g). HCA7 cells showed a similar effect to SW620 cells, however LS174T cells did not show significantly increased sensitivity to UK-5099 with long-term aspirin (Supplementary Figure 4a-b), suggesting some cell-line specificity in this response. This effect was also investigated upon short-term aspirin treatment in SW620 cells (Supplementary Figure 4c), which also increased sensitivity to UK-5099.

These findings support the hypothesis that when treated with aspirin, cells reprogramme their metabolism in order to maintain proliferation (summarised in Figure 5a), leaving them vulnerable to further metabolic manipulation. While they have sufficient metabolic plasticity to prevent an impact on ATP production and complete inhibition of proliferation in the presence of aspirin alone, the cells become more reliant on particular metabolic pathways and are left vulnerable to their targeting, leading to further impaired proliferation and cell death.

**Figure 5.**
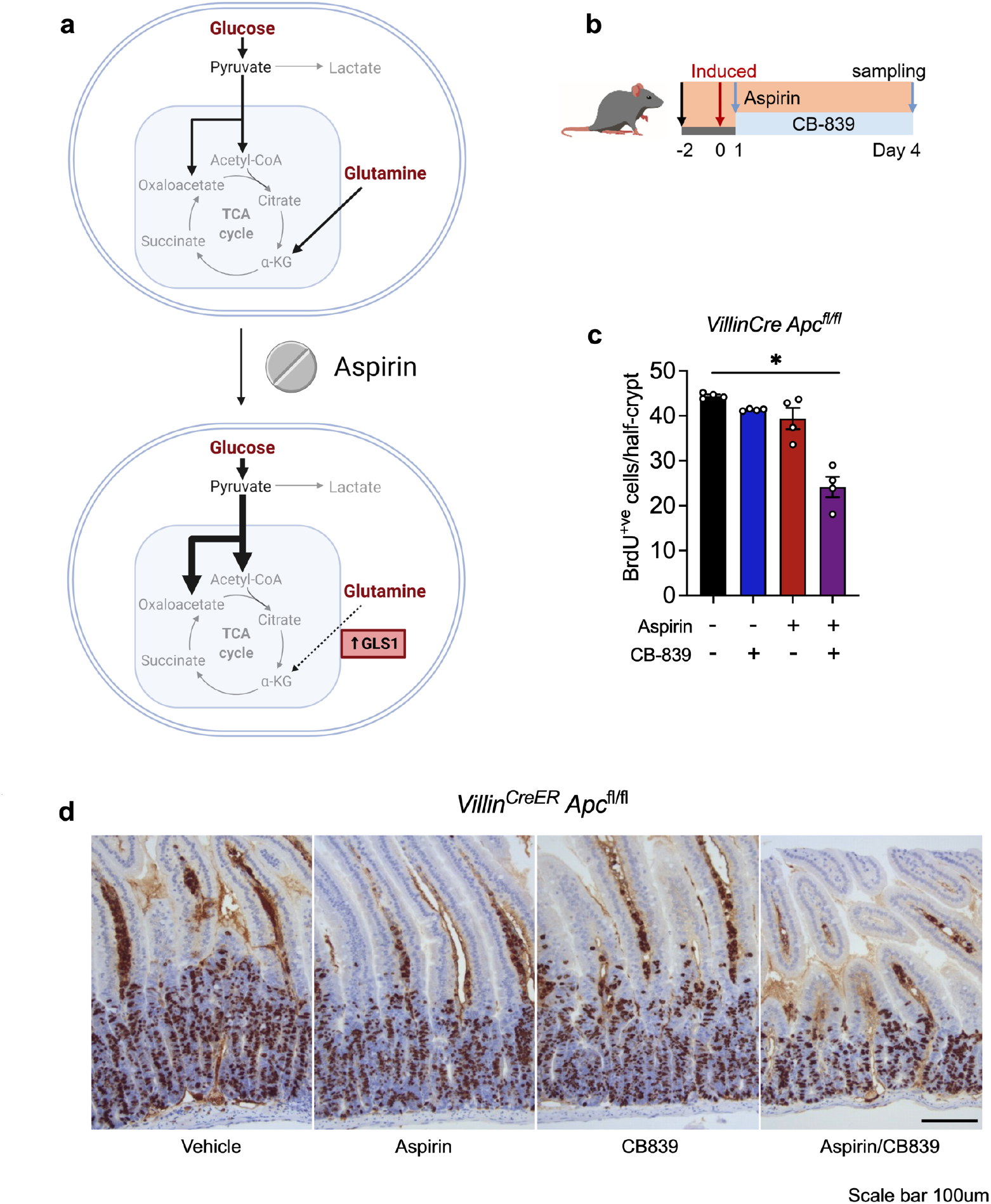
Aspirin and CB-389 in combination reduce colon crypt proliferation *in vivo*. **a)**Schematic summarising the effects of aspirin treatment on metabolic reprogramming of CRC cells. Created with BioRender.com. **b)** Schematic showing the timeline of treatment, induction with tamoxifen (2 mg) and sampling for *in vivo* aspirin (2.6mg/ml in drinking water) and CB-839 (200mg/kg) combination experiments using the *Villin^CreER^ Apc^fl/fl^* mouse model. **c-d)** Quantification of BrdU and representative images of BrdU staining in small intestine of *Villin^CreER^ Apc^fl/fl^* mice treated with vehicle, CB-839 (200mg/kg) and/or aspirin (2.6mg/ml in drinking water). Scale bar 100μm. Error bars show SEM (n=4 mice per experimental arm). Each dot represents the average number of BrdU positive cells per half crypt for each mouse. Asterisks refer to p-values obtained from one-tailed Mann-Whitney tests (*=p<0.05; **=p<0.01).

### Aspirin and CB-389 in combination reduce colon crypt proliferation *in vivo*

We next sought to investigate the efficacy of combining aspirin and CB-839 in vivo. To achieve this, we used the well characterized VillinCreER Apcfl/fl mouse model. The mice were induced with tamoxifen (2mg on two consecutive days) to conditionally delete Apc throughout the intestinal epithelium. The mice were treated with either vehicle or aspirin (2.6mg/ml) or CB-839 (200mg/kg) alone or in combination (aspirin + CB-839) and the effect on intestinal epithelial cell proliferation was investigated (treatments summarised in Figure 5b). Apc loss leads to a characteristic crypt hyperproliferation in this model as assessed by BrdU incorporation and quantification of the number of stained BrdU+ cells in each crypt (Figure 5c-d). Strikingly, the combination of aspirin (2.6mg/ml) and CB-839 (200mg/kg) led to a significant suppression of crypt hyperproliferation in the small intestine as indicated by BrdU stained cells, while neither aspirin nor CB-839 alone impacted proliferation (Figure 5c-d). These results support our *in vitro* findings showing that aspirin induces sensitivity to CB-839 and support the potential for clinical utility of this approach.

## Discussion

Further understanding of the anti-cancer cellular mechanisms of aspirin will be beneficial to inform patient stratification and maximise its benefits. Although best known as a cyclooxygenase (COX) inhibitor, aspirin is a highly pleiotropic drug with many cellular targets. Despite the importance of COX/PGE2 signalling in cancer (28), this mechanism is not sufficient to fully explain the anti-cancer effects of aspirin (29). Studies have shown aspirin treatment impacts many other pathways including Wnt (30), NF-κB (31, 32), AMPK and mTOR (33, 34), as well as causing epigenetic alterations such as histone methylation (35, 36), however, relatively little work has focused on the impact of aspirin treatment on cancer cell metabolism.

Here, we used proteomics with the aim to identify novel cellular mechanisms of aspirin that may contribute to its anti-cancer effect. This analysis highlighted a potential effect on cellular metabolism which was subsequently confirmed; using a combination of extracellular flux analysis and SIL we comprehensively characterised the effect of aspirin on CRC cell metabolic pathway activity. Our results show that although there is no overall impact of aspirin on ATP production, long-term aspirin exposure leads to significant reprogramming of nutrient utilisation, leading to a reduction in glutaminolysis and increased glucose entry into the TCA cycle in three CRC cell lines (summarised in Figure 5a).

We also show regulation of proteins involved in central carbon metabolism that is consistent with the effects on pathway activity, including increased expression of PC and decreased expression of PDK1 and regulation of several glutaminolysis enzymes. Surprisingly, there was a strong increase in expression of GLS1, which is contradictory to the observed decrease in net glutaminolysis. We suggest that this is a compensatory response to glutaminolysis being otherwise impaired. Increased glucose utilisation is also a likely compensatory mechanism to reduced glutaminolysis, as both nutrients provide important carbon sources for the TCA cycle. This information was used to infer potential metabolic vulnerabilities induced by aspirin; we show that aspirin treated cells are more sensitive to the GLS1 inhibitor CB-839 and MPC1 inhibitor UK-5099. Importantly, the combination of aspirin and CB-839 was found to be effective in reducing cell proliferation both *in vitro* and *in vivo*; the treatment inhibited proliferation and induced apoptosis in CRC cell lines, and inhibited proliferation of colonic epithelial cells in *VillinCre;Apc^fl/fl^* mice.

A small number of recent studies have linked aspirin’s anti-cancer effects to metabolism; aspirin has been found to inhibit glucose metabolism through regulation of PDK1 in breast cancer cells (37), and through regulation of GLUT1 in hepatoma cells (38). In support of our study, Boku et al., show that combining GLS1 inhibition with aspirin is effective at reducing colony-forming efficiency of CRC cells *in vitro* (39). The same study suggested that aspirin treatment mimics the effects of glutamine deficiency in CRC cells, and showed increased expression of glutaminolysis genes with aspirin treatment (39). The authors concluded that glutaminolysis is therefore likely upregulated upon aspirin treatment, however this was not directly measured. Interestingly, our functional studies of glutaminolysis using SIL show a reduction in glutaminolysis with aspirin. We conclude that the upregulation of GLS1 is a likely compensatory mechanism for reduced glutaminolysis. Despite these apparent differences, both studies highlight the possibility that aspirin has potential to be a simple and cost-effective drug to increase the efficacy of CB-839 in CRC patients. This is important as despite the attractiveness of targeting glutamine metabolism for cancer therapy (40), CB-839 has achieved varying success in previous studies, particularly when studied *in vivo* (41, 42). Indeed, in this study, CB-839 alone was almost completely ineffective at inhibiting cell proliferation and had no effect on apoptosis *in vitro* or crypt proliferation *in vivo*. Therefore, our findings have exciting implications for clinical translation, as both drugs are already used clinically and have known safety profiles in humans.

An increasing number of studies highlight the value of combinatory approaches when applying metabolic interventions. One comprehensive study highlights several combinations of metabolic inhibition that overcome the ability of cancer cells to adapt their metabolism in response to singular perturbations – known as metabolic flexibility or plasticity (19). A recent study showed that combining CB-839 with an inhibitor of ASCT2 was effective in liver cancer cells (43). Several studies have also highlighted success when combining metabolic interventions with chemotherapies (44–46) and immunotherapies (47). There is also increasing interest in the impact of diet on tumour metabolism and how this may interact with metabolic therapies (48). Our findings support the value of combining multiple complementary interventions when targeting tumour metabolism. By highlighting metabolic vulnerabilities caused by the exposure of CRC to aspirin alone, we were able to exploit these to maximise inhibition of CRC cell proliferation with metabolic inhibitors. The use of aspirin to increase efficacy of metabolic inhibitors is particularly valuable due to the longstanding ubiquitous use of aspirin in clinical practice. The efficacy of aspirin and CB-839 *in vivo* shown here suggest that this combination warrants further clinical studies and could potentially provide a novel therapeutic option for CRC patients alongside traditional chemotherapies.

## Acknowledgements

We would like to thank D. Avizonis and L. Choinière from the McGill University Metabolomics Core Facility for their support. We would also like to thank P. Lewis and K. Heesom in the Proteomics Facility at the University of Bristol.

AKH and AJH were supported by the James Tudor Foundation, AKH, AJH and EMHM were supported by John and Bridget Maynard, PT was supported by a PhD studentship from Bowel & Cancer Research. AKH, AJH, EMHM and PT were also supported by the John James Bristol Foundation. DF, TJC, OJS and ACW by an MRC Research Grant (MR/R017247/1). EEV and DNL are supported by Diabetes UK (17/0005587) and the World Cancer Research Fund (WCRF UK), as part of the World Cancer Research Fund International grant program (IIG_2019_2009). EEV and TJC are supported by the Cancer Research UK (CRUK) Integrative Cancer Epidemiology Programme (C18281/A29019). AKN was supported by Cancer Research UK (CRUK) Beatson Institute core funding (A17196, A31287) - awarded to OJS. OJS was supported by CRUK grants (A21139, A12481, A17196 and A31287) and ERC Starting grant (311301).

## Methods

### Cell lines and culture

The human colorectal carcinoma-derived cell lines; SW620, LS174T were obtained from the American Type Culture Collection (ATCC, Maryland, USA) and HCA7 was a kind gift from Dr Susan Kirkland, Imperial College, London. All cell lines were routinely tested for mycoplasma contamination using MycoAlert PLUS mycoplasma detection kit (Lonza, MD, USA) and molecularly characterised using an “in house” panel of cellular and molecular markers to check that cell lines have not been cross contaminated (every 3-6 month). Stocks were securely catalogued and stored; passage numbers strictly adhered to preventing phenotypic drift. All cell lines were cultured in Dulbecco’s Modified Eagle Medium (DMEM) (Sigma-Aldrich, Merck, KGAa) with added 10% foetal bovine serum (FBS) (Sigma-Aldrich, Merck, KGaA), 2 mM glutamine (Gibco, ThermoFisher Scientific Inc.), 100 units/ml penicillin and 100 units/ml streptomycin (Gibco, ThermoFisher Scientific Inc). For stock purposes, cells were maintained in 25cm^2^ tissue culture (T25) flasks (Corning, NY, USA) and incubated at 37°C in dry incubators maintained at 5% CO^2^. Cell media were changed every 3-4 days. Experiments were performed in triplicate independently with distinct passages of cells, unless otherwise states.

### Treatments

#### Long-term aspirin

For the long-term aspirin treated cells, a 20mM stock solution of aspirin (Sigma, Merck KGaA, Darmstadt, Germany) was created by adding 3.6mg/ml aspirin to 10% DMEM, fresh aspirin was made up immediately prior to use. Aspirin concentration in the growth media was maintained continuously for ~52 weeks. Passage frequency and ratio was adjusted to maintain confluency in aspirin treated cells.

#### CB-839 and UK-5099

Stock of CB-839 (Sigma-Aldrich, Merck, KGaA) and UK-5099 (Sigma Aldrich, Merck, KGaA) were made in DMSO. DMSO concentration was maintained consistently in all treatment concentrations and vehicle control.

### Proliferation assays

#### Crystal violet staining

To measure proliferation of cells treated with aspirin in combination with either CB-839 or UK-5099, cells were seeded into 96 well plates (Corning, NY, USA) (20,000 cells per well in all conditions except HCA7 cells treated with 4mM aspirin, where 40,000 cells per well were seeded) in normal growth medium and incubated for 24 hours, with 3-4 technical replicate wells per treatment condition. Cells were then treated with media containing drug treatments (or vehicle control) and incubated for a further 72 hours. Plates were then fixed with 4% PFA for 15 minutes, then stained with 0.5% crystal violet solution (Sigma-Aldrich, Merck KGaA), before solubilisation in 2% SDS and subsequent OD595 measurements were obtained using an iMark microplate reader (Bio-Rad, Laboratories, Inc.). The number of adherent cells was claculated by the confluence in each concentration of CB-839/UK-5099 at 72 hours, relative to the same concentration of aspirin in control conditions, in order to compare the effect of the drugs between different aspirin treatments.

For experiments using Human Plasma-Like Medium (HPLM – Gibco, ThermoFisher Scientific, Inc.), HPLM was supplemented with 10% dialysed FBS (dFBS) and experiments were performed as above. Cells were incubated in 10% dFBS HPLM for at least 48 hours prior to the start of the treatment, to allow for metabolic adaptation. During the experiment, media was changed every 24 hours (unlike experiments in DMEM where the same media was left for the full 72 hours of treatment), in order to avoid depletion of the low levels of nutrients.

#### IncuCyte

To simultaneously measure cell proliferation and apoptosis upon treatment with aspirin in combination with CB-839, a IncuCyte ZOOM live cell imaging system was used. Cells were seeded in 96 well plates (20,000 cells per well) and incubated for 24 hours, with 3-4 wells per treatment condition (technical replicates). Cells were then treated with treatment-containing media (or vehicle control) and placed in the IncuCyte system. Percentage of Confluence was measured every four hours for the total time indicated on the graphs. The IncuCyte system took four different image fields per well. At time of treatment the cells were also treated with 2μM CellEvent caspase-3/7 green detection reagent (C10423; Invitrogen, ThermoFisher Scientific, Inc), which was used to measure apoptosis. Green fluorescent cells, indicating active caspase-3/7 and apoptosis, were measured by the IncuCyte system as green object count (1/mm^2^). For each individual well, green object count was normalised to confluence at each timepoint, and results expressed as relative apoptosis. The same method was performed using SW620 cells treated with 2μM ABT-737 compared to a vehicle control prior to the start of the assay, as a positive control for apoptosis in order to validate this assay and confirm detection of apoptotic cells.

### Proteomic analysis

#### TMT labelling and high pH reversed-phase chromatography

Following 52 weeks of aspirin treatment to develop long-term treated cell lines, cells were seeded in T25 flasks, and following maintenance in aspirin for a further 72 hours whole cell protein lysates were collected. Lysates were collected as described previously (49). Protein concentrations were ascertained, and samples adjusted to 2 mg/mL. 100ug of each sample was digested with trypsin overnight at 37°C, labelled with Tandem Mass Tag (TMT) ten plex reagents according to the manufacturer’s protocol (ThermoFisher Scientific, Inc) and the labelled samples pooled.

A aliquot of the pooled sample was evaporated to dryness, resuspended in 5% formic acid and then desalted using a SepPak cartridge according to the manufacturer’s instructions (Waters, Milford, Massachusetts, USA). Eluate from the SepPak cartridge was again evaporated to dryness and resuspended in buffer A (20 mM ammonium hydroxide, pH 10) prior to fractionation by high pH reversed-phase chromatography using an Ultimate 3000 liquid chromatography system (ThermoFisher Scientific, Inc). In brief, the sample was loaded onto a XBridge BEH C18 Column (130Å, 3.5 μm, 2.1 mm × 150 mm, Waters, Milford, Massachusetts, USA) in buffer A and peptides eluted with an increasing gradient of buffer B (20 mM Ammonium Hydroxide in acetonitrile, pH 10) from 0-95% over 60 minutes. The resulting fractions (15 in total) were evaporated to dryness and resuspended in 1% formic acid prior to analysis by nano-LC MSMS using an Orbitrap Fusion Tribrid mass spectrometer (ThermoFisher Scientific, Inc).

#### Nano-LC Mass Spectrometry

High pH RP fractions were further fractionated using an Ultimate 3000 nano-LC system in line with an Orbitrap Fusion Tribrid mass spectrometer (ThermoFisher Scientific, Inc). In brief, peptides in 1% (v/v) formic acid were injected onto an Acclaim PepMap C18 nano-trap column (ThermoFisher Scientific, Inc). After washing with 0.5% (v/v) acetonitrile 0.1% (v/v) formic acid peptides were resolved on a 250 mm × 75 μm Acclaim PepMap C18 reverse phase analytical column (ThermoFisher Scientific, Inc) over a 150 min organic gradient, using 7 gradient segments (1-6% solvent B over 1 min., 6-15% B over 58 min, 15-32% B over 58min., 32-40%B over 5min., 40-90%B over 1min., held at 90%B for 6min and then reduced to 1%B over 1min.) with a flow rate of 300nl min-1. Solvent A was 0.1% formic acid and Solvent B was aqueous 80% acetonitrile in 0.1% formic acid. Peptides were ionized by nano-electrospray ionization at 2.0kV using a stainless-steel emitter with an internal diameter of 30μm (Thermo Scientific) and a capillary temperature of 275°C.

All spectra were acquired using an Orbitrap Fusion Tribrid mass spectrometer controlled by Xcalibur 2.1 software (Thermo Scientific) and operated in data-dependent acquisition mode using an SPS-MS3 workflow. FTMS1 spectra were collected at a resolution of 120 000, with an automatic gain control (AGC) target of 200 000 and a max injection time of 50ms. Precursors were filtered with an intensity threshold of 5000, according to charge state (to include charge states 2-7) and with monoisotopic peak determination set to peptide. Previously interrogated precursors were excluded using a dynamic window (60s +/−10ppm). The MS2 precursors were isolated with a quadrupole isolation window of 1.2m/z. ITMS2 spectra were collected with an AGC target of 10 000, max injection time of 70ms and CID collision energy of 35%.

For FTMS3 analysis, the Orbitrap was operated at 50 000 resolution with an AGC target of 50 000 and a max injection time of 105ms. Precursors were fragmented by high energy collision dissociation (HCD) at a normalised collision energy of 60% to ensure maximal TMT reporter ion yield. Synchronous Precursor Selection (SPS) was enabled to include up to 5 MS2 fragment ions in the FTMS3 scan.

#### Data Analysis

The raw data files were processed and quantified using Proteome Discoverer software v2.1 (ThermoFisher Scientific, Inc) and searched against the UniProt Human database (downloaded September 2017; 140000 sequences) using the SEQUEST HT algorithm. Peptide precursor mass tolerance was set at 10ppm, and MS/MS tolerance was set at 0.6Da. Search criteria included oxidation of methionine (+15.995Da), acetylation of the protein N-terminus (+42.011Da) and methionine loss plus acetylation of the protein N-terminus (−89.03Da) as variable modifications and carbamidomethylation of cysteine (+57.021Da) and the addition of the TMT mass tag (+229.163Da) to peptide N-termini and lysine as fixed modifications. Searches were performed with full tryptic digestion and a maximum of two missed cleavages were allowed. The reverse database search option was enabled and all data was filtered to satisfy false discovery rate (FDR) of 5%.

#### Protein abundance processing

Protein groupings were determined by PD2.1, however, the master protein selection was improved with an in-house script. This enables us to infer biological trends more effectively in the dataset without any loss in the quality of identification or quantification. The MS data were searched against the human Uniprot database retrieved on 2019-10-02, and updated with additional annotation information on 2020-04-21.

The protein abundances were normalised within each sample to total peptide amount, then Log2 transformed to bring them closer to a normal distribution.

#### Statistics

Statistical significance was then determined using a Welch’s T-Tests between the conditions of interest. The p-values were FDR corrected using the Benjamini-Hochberg method.

#### QIAGEN Ingenuity Pathway Analysis (QIAGEN IPA)

Data were analysed using Ingenuity Pathway Analysis. Proteins from the dataset that met the cut-off of p<0.05 were considered for the analysis, and compared a reference set consisting of the full list of proteins identified in the experiment. A right-tailed Fisher’s Exact Test was used to calculate a p-value determining the probability that the association between the genes in the dataset and the pathways/upstream regulators/functions were by chance alone, and the predicted and observed regulation patterns of the proteins were used to predict an activation z-score.

#### Overrepresentation analysis

Overrepresentation analysis was performed using Webgestalt (www.webgestalt.org), using the functional databases Geneontology, and pathways (KEGG). Gene symbols were entered for proteins that showed signification regulation (p<0.05, fold change>1.4 or <0.71), in both 2mM and 4mM long-term aspirin compared to control.

### SDS-PAGE and immunoblotting

Cell lysates were prepared and subjected to western analysis as described previously (49) using antibodies to the following: α-tubulin (T9026, Sigma-Aldrich, Merck, Inc.), ATF4 (11815, Cell Signaling Technology Inc. (CST)), ASCT2 (5345, CST), GLS1 (88964, CST), GPT2 (16757-1-AP, ProteinTech, Group, Inc.), LAT1 (5347, CST), PC (ab126707, Abcam, Cambridge, UK), PDK1 (3820, CST). All were used at 1:1000 dilution, except for α-tubulin which was used at 1:10,000 dilution. Density of bands detected by immunoblotting was measured using ImageJ. In order to compare data from independent experiments, protein expression changes were calculated as relative changes from the control conditions in each replicate. Results from at least three independent experiments were analysed with this method.

### Quantitative reverse transcriptase-PCR (qPCR)

Total RNA was extracted using Tri-Reagent (Sigma-Aldrich) as per manufacturer’s instruction an RNAeasy mini kit with an ON-column DNA digest step (Qiagen, Limberg, Netherlands) was used according to the manufacturer’s instructions to clean up the RNA. RNA concentration and purity was measured using a NanoDrop™ spectrophotometer (ThermoFisher Scientific Inc.). Synthesis of cDNA and qRT-PCR were performed as previously described (50) using the following primers (all from Qiagen, Limberg, Netherlands): *ATF4* (QT00074466), *GPT2* (QT00066381), *GLS1* (QT00019397), *PC* (QT01005592), *PDK1* (QT00069636), *SLC7A11* (QT00002674), *SLC7A5* (QT00089145). Gene expression was normalised to the housekeeping genes *TBP* (cat. no. QT00000721) or *HPRT* (QT00059066), both from Qiagen.

### Extracellular flux analysis

Extracellular flux analysis was carried out using the XFp Seahorse Extracellular Flux Analyzer (Agilent), according to the manufacturer’s protocol. Long-term aspirin treated SW620 cells (60,000 cells) were seeded onto a Cell-Tak (354240, Corning, NY, USA)-coated microplate (2-3 technical replicate wells per condition) and centrifuged at 200g for 1 minute (no brake), allowing for immediate adhesion. Cells were seeded in Seahorse XF assay media (Agilent, CA, USA) supplemented with 10mM glucose, 2mM glutamine and 1mM pyruvate (Agilent, CA, USA). Corresponding OCR/ECAR (oxygen consumption rate/extracellular acidification rate) changes were monitored for the duration of the experiment. Wells had subsequent injections of oligomycin (2μM), FCCP (2μM), antimycin A (1μM) and rotenone (1μM) and monensin (20μM) (in order to determine maximal glycolytic rate, as shown by Mookerjee et al. (51, 52)) all from Sigma-Aldrich. Data were acquired using the Seahorse Wave software v2.6 (Agilent, CA, USA). The experiment was performed independently in triplicate.

### Stable isotope tracer analysis

For stable isotope labelling (SIL) experiments, cells were cultured with U-[^13^C]-Glc or U-[^13^C]-Q (Cambridge Isotopes Laboratories, Inc.) for the indicated time points. ^13^C labelled nutrients were added to glucose, glutamine free DMEM, supplemented with 10% dFBS, 100 units/ml penicillin and 100 units/ml streptomycin, 10mM glucose and 2mM glutamine (^13^C labelled or unlabelled as appropriate). Cellular metabolites were extracted and analysed by gas chromatography-mass spectrometry (GC-MS) using protocols described previously (53–55). Metabolite extracts were derived using N-(tert-butyldimethylsilyl)-N-methyltrifluoroacetamide (MTBSTFA) as described previously (56). D-myristic acid (750ng/sample) was added as an internal standard to metabolite extracts, and metabolite abundance was expressed relative to the internal standard and normalized to cell number. Mass isotopomer distribution was determined using a custom algorithm developed at McGill University (55). The experiment was performed with three different flasks of cells from the same passage number per condition.

### *In vivo* experiments

All *in vivo* experiments were carried out in accordance with the UK Home Office regulations (under project licences: 70/8646 and PP3908577), and by adhering to the ARRIVE guidelines with approval from the Animal Welfare and Ethical Review Board of the University of Glasgow. Mice were housed under a 12 h light-dark cycle, at constant temperature (19-23 °C) and humidity (55 ± 10%). Standard diet and water were available *ad libitum*. The majority of the work was performed in the C57BL/6J background. The following alleles were used in this study: *VillinCreER* (57), *Apc^fl^* (58). Full intestinal recombination was obtained by two intraperitoneal injections of 2mg tamoxifen and tissues were harvested 4 days post induction. For drug studies *in vivo*, Villin-Cre^ERT2^ *Apc^fl/fl^* mice were treated with CB-839 (200mg/kg in 25% (w/v) hydroxypropyl-β-cyclodextrin in 10mm citrate at pH 2.0) or vehicle from day 1 post i.p. tamoxifen administration. For aspirin and combination treatments, mice received aspirin (2.6mg/ml in drinking water) two days prior to tamoxifen administration and remained on aspirin till the end of the study. Animals were injected with BrdU (i.p.) 2 hours prior to sampling tissues.

### Immunohistochemistry (IHC)

Mouse intestines were flushed with water, cut open longitudinally, pinned out onto silicone plates and fixed in 10% neutral buffered formalin overnight at 4°C. Fixed tissue was rolled from proximal to distal end into swiss-rolls and processed for paraffin embedding. Tissue blocks were cut into 5μm sections and stained with haematoxylin and eosin (H&E). IHC was performed on formalin-fixed intestinal sections according to standard staining protocols. Primary antibody used was against BrdU (1:150, BD Biosciences, #347580), representative images are shown.

### Statistical analysis

Data were presented and statistical analysis was performed using GraphPad Prism 9. Statistical tests were performed as stated, and significance was expressed as *=p<0.05, **=p<0.01, ***=p<0.001, ****=p<0.0001. Results are expressed as mean values with standard error of the mean (SEM) where independent experiments are compared, and with standard deviation (SD) where technical replicates compared. Here, technical replicates refer to separate wells or flasks of cells that are from the same original passage of cells, seeded and treated at the same time. Independent experiments refer to separate passages of cells that were seeded at different times.

## Notes

### Competing Interest Statement

The authors have declared no competing interest.

### Summary of Updates

Figure 2 updated for visual clarity

