## Supplementary Material for "Aspirin reprogrammes colorectal cancer cell metabolism and sensitises to glutaminase inhibition"

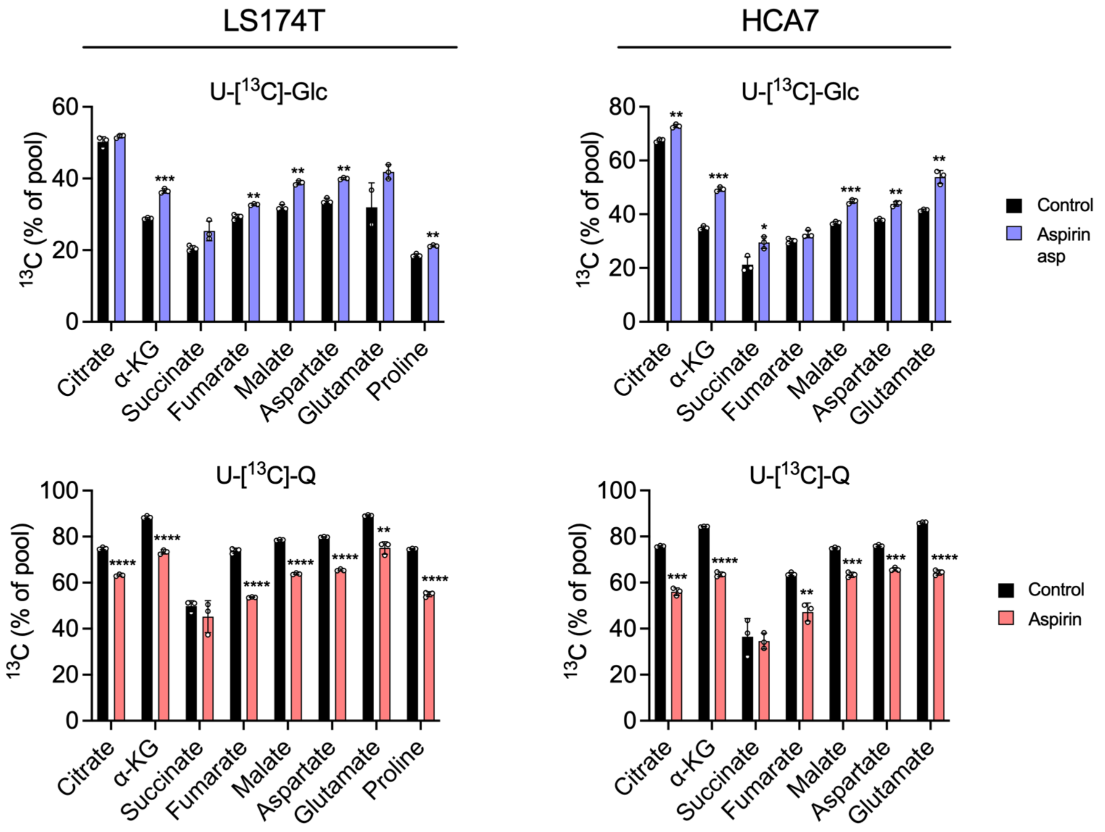


**Supplementary Figure 1. Aspirin treatment reprogrammes nutrient utilisation in CRC cells.** Proportion of $^{13}C$ labelling in downstream metabolites after 8 hours incubation with either U-[$^{13}C$]-Glc or U-[$^{13}C$]-Q in LS174T and HCA7 cells with long-term 4mM aspirin treatment compared to control cells. Error bars represent SD (n=3 technical replicates). Asterisks refer to adjusted p-values obtained from multiple t tests (*=p<0.05, **=p<0.01, ***=p<0.001, ****=p<0.0001).

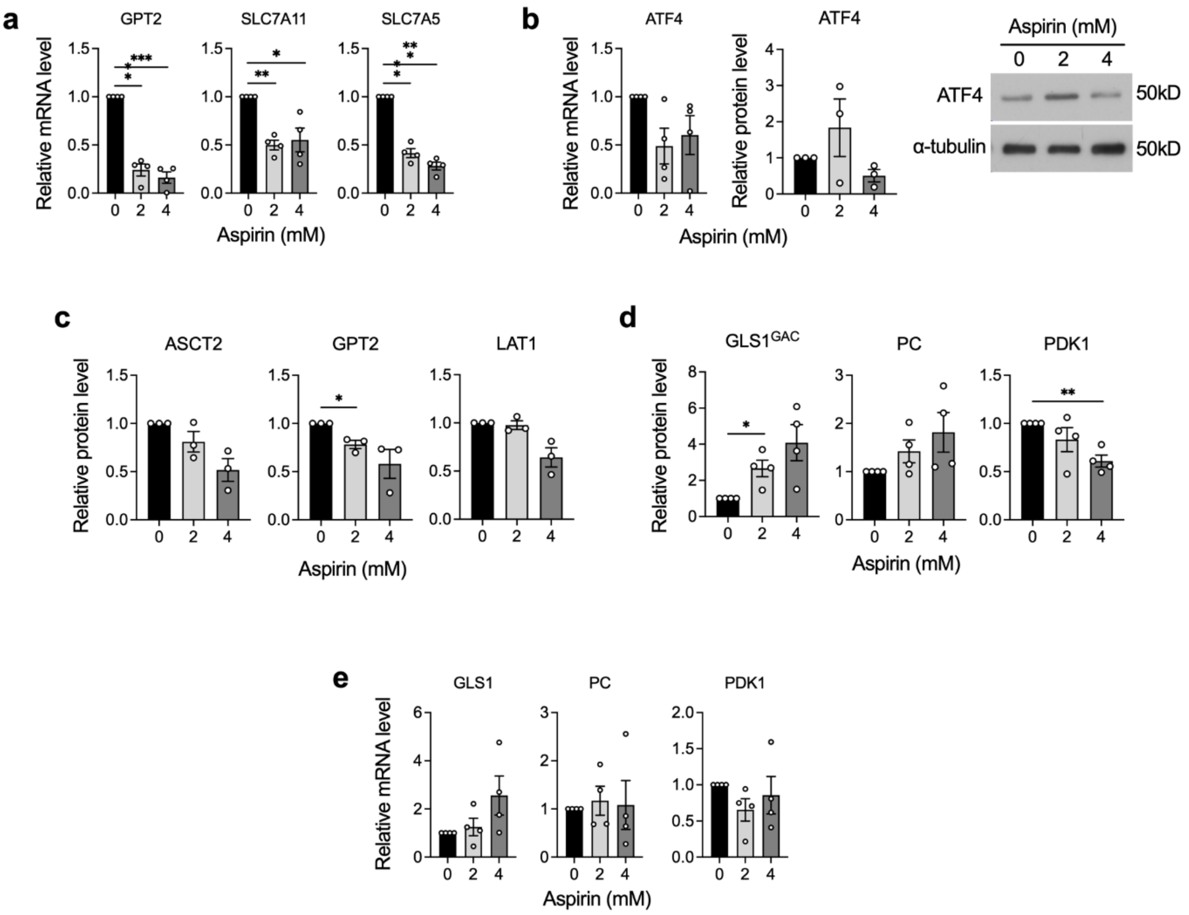


**Supplementary Figure 2. Long-term aspirin treatment regulates metabolic enzyme expression in SW620 cells. a-e)** qPCR analysis and quantification of immunoblot of metabolic gene expression in long-term (52 week) aspirin exposed SW620 cells. Error bars represent SEM (n≥3 independent experiments). Asterisks refer to p-values obtained from one-sample t tests, comparing to a hypothetical mean of 1 (*=p<0.05, **=p<0.01, ***=p<0.001). **a**) qPCR analysis of ATF4 target gene expression. **b)** qPCR and quantified immunoblot analysis of ATF4 expression. Immunoblot image shows a representative blot of 3 independent experiments, α-tubulin is used as a loading control. **c)** Quantification of immunoblots for proteins involved in glutamine metabolism. **d)** Quantification of immunoblot for proteins involved in central carbon metabolism. $GLS1^{GAC}$ = GAC splice isoform of glutaminase 1. **e)** Relative mRNA levels, determined by qPCR of genes involved in central carbon metabolism.

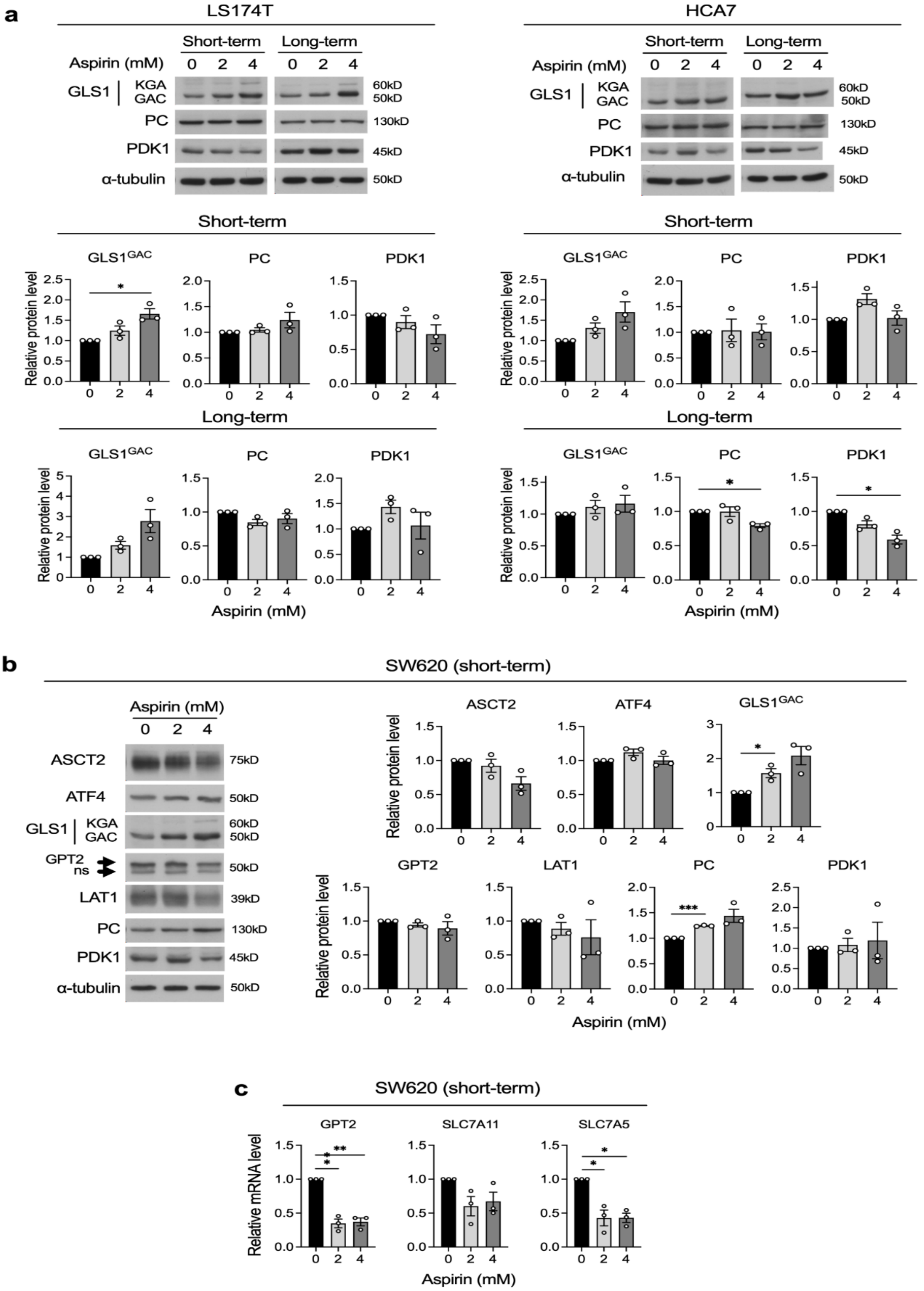


**Supplementary Figure 3. Long-term and short-term aspirin treatment regulates metabolic enzyme expression in three CRC cell lines. a-b)** Graphs show quantification of immunoblotting, error bars represent SEM (n=3 independent experiments). Asterisks refer to p-values obtained from one-sample t tests, comparing to a hypothetical mean of 1 (*=p<0.05, ***=p<0.001) Immunoblot images are representative of three independent experiments, α-tubulin is used as a loading control. $GLS1^{GAC}$ = GAC splice isoform of glutaminase 1. **a)** Short-term (72 hour) and long-term aspirin (52 week) treated LS174T and HCA7 cells. **b)** Short-term (72 hour) aspirin treated SW620 cells. **c)** qPCR analysis of ATF4 target genes in short-term (72 hour) aspirin treated SW620 cells. Error bars represent SEM (n=3 independent experiments), asterisks refer to p-values obtained from one-sample t tests, comparing to a hypothetical mean of 1 (*=p<0.05, **=p<0.01).

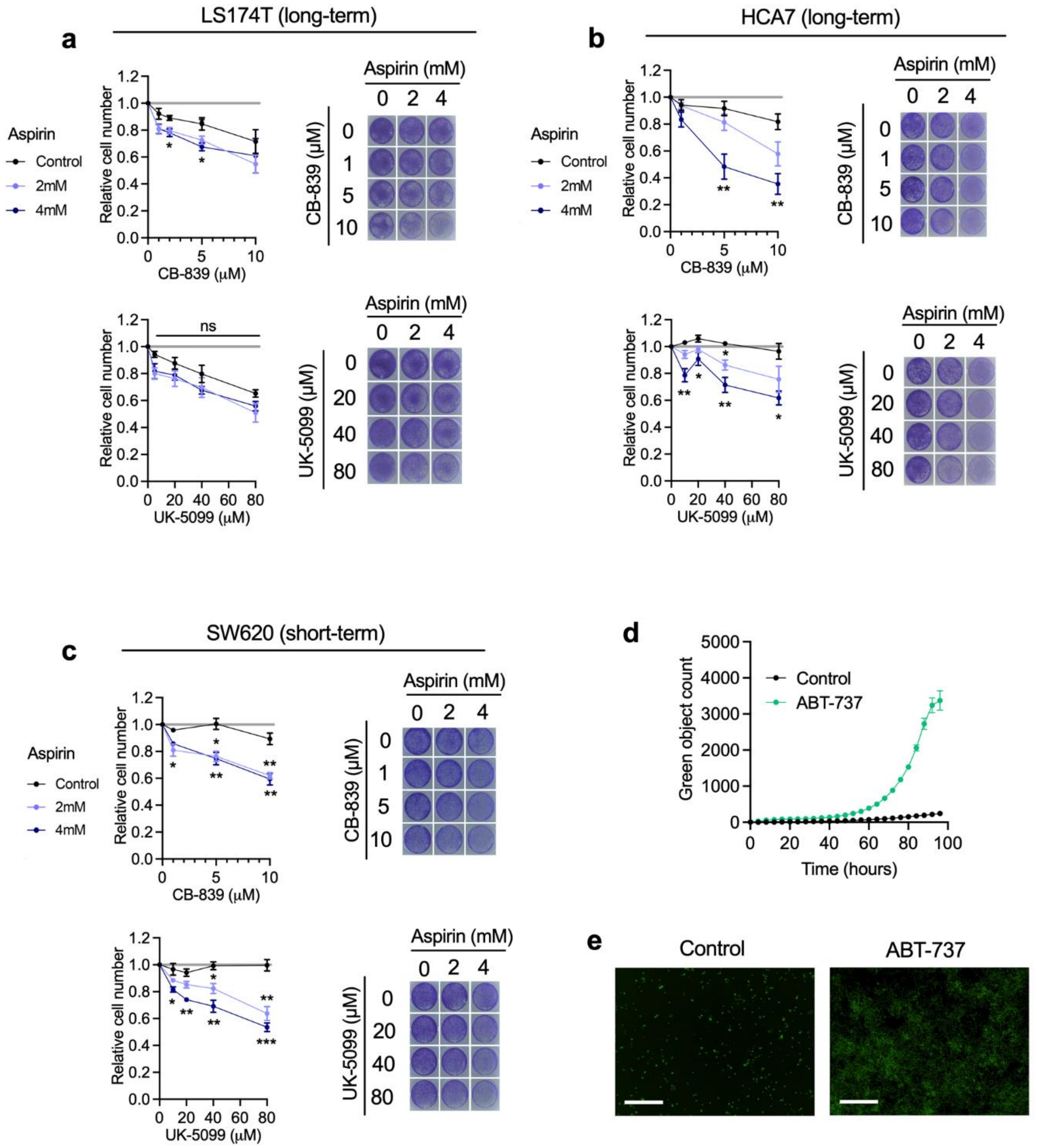


**Supplementary Figure 4. Aspirin sensitises CRC cells to metabolic inhibitors. a-c)** Cell proliferation assays of long-term (52 week) aspirin treated LS174T cells **(a)** and HCA7 cells **(b)** and short-term (72 hour) aspirin treated SW620 cells **(c)** with increasing concentrations of either CB-839 or UK-5099. Graphs show relative cell number in each aspirin condition as measured by crystal violet staining at 72 hours of drug treatment compared to vehicle control. Error bars show SEM (n=5 independent experiments for HCA7 with CB-839, n=3 independent experiments for all other experiments). Asterisks refer to p-values obtained using one-way ANOVAs with Dunnett's multiple comparisons tests at each drug concentration (*=p<0.05, **=p<0.01, ***=p<0.001). Images show representative wells in each condition after 72 hours of treatment. **d)** Quantification of green fluorescent nuclei indicating apoptotic SW620 cells with activated caspase-3/7, with 2µM ABT-737 treatment compared to control, as a positive control for apoptosis. Error bars represent SD (n=3 technical replicates). **e)** Representative images of 2µM ABT-737 treated and control SW620 cells at the end of the assay (~96 hours). Scale bar represents 300µm.
